# Generation of Human Regulatory Dendritic Cells from Cryopreserved Healthy Donor Cells and Hematopoietic Stem Cell Transplant Recipients

**DOI:** 10.1101/2023.08.11.552986

**Authors:** Sabrina M. Scroggins, Annette J. Schlueter

**Affiliations:** Deparment of Biomedical Sciences, University of Minnesota-Duluth, Duluth, Minnesota 55812; Deparment of Obstetrics and Gynecology; Department of Pathology, University of Iowa, Iowa City, Iowa 52242

**Author notes:** **Corresponding author** Sabrina M. Scroggins, PhD, University of Minnesota-Duluth, 1035 University Drive, 341 SMED, Duluth, MN 55812.

**Keywords:** good manufacturing practices (GMPs), graft versus host disease (GVHD), hematopoietic stem-cell transplantation (HSCT), monocytes, regulatory dendritic cells (DCreg), human

## Abstract

Acute graft versus host disease (GVHD) remains a significant complication following hematopoietic stem cell transplant (HSCT), despite improved human leukocyte antigen (HLA) matching and advances in prophylactic treatment regimens. Previous studies have shown promising results for future regulatory dendritic cell (DCreg) therapies in the amelioration of GVHD. This study evaluates the effects of cryopreservation on DCreg generation, generation of young and older DCreg in serum-free media, and the feasibility of DCreg generated from young and older HSCT donor monocytes. DCreg were generated in X-vivo 15 serum-free media from donor monocytes. Donors included young and older individuals, either healthy donors or HSCT patients. Phenotypic differences in cell populations were assessed via flow cytometry while pro-inflammatory and anti-inflammatory cytokine production was evaluated in culture supernatants. The number of DCreg generated from cryopreserved monocytes of healthy donors was not significantly different from freshly isolated monocytes. DCreg generated from cryopreserved monocytes had similar levels of co-stimulatory molecule expression, inhibitory molecule expression, and cytokine production as freshly isolated monocytes. Young and older healthy donor monocytes generated similar numbers of DCreg with similar cytokine production and phenotype. Although monocytes from older HSCT patients produced significantly fewer DCreg, DCreg from young and older HSCT patients have a comparable phenotype and cytokine production. Monocytes from young and older myelodysplastic syndrome (MDS) patients generated reduced numbers of DCreg compared to non-MDS monocytes. Results suggest cryopreservation of monocytes from many HSCT patients allows for cost effective generation of DCreg for the prevention and treatment of GVHD on an as needed basis. Although generation of DCreg from MDS patients require further assessment, these data support the possibility of *in vitro* generated DCreg as a therapy to reduce GVHD-associated morbidity and morbidity in young and older HSCT recipients.

## Introduction

Acute graft versus host disease (GVHD) is a systemic immune mediated disease primarily involving the skin, liver, and gastrointestinal tract ^1-4^. Despite improved human leukocyte antigen (HLA) matching and advances in prophylactic treatment regimens, GVHD following hematopoietic stem cell transplant (HSCT) remains a significant complication. GVHD occurs in 40-85% of young and up to 70% of older allogeneic transplant recipients and is fatal in 30-45% of those patients ^5-8^. To this end, age is an independent risk factor for increased incidence and severity of GVHD ^4,9-11^ and the number of older patients undergoing transplant each year are steadily increasing ^12^. As the total number of HSCT performed each year is steadily increasing, so is need for alternative therapies for GVHD in both young and older HSCT recipients.

In a murine model of GVHD, a single treatment with age-matched, syngeneic regulatory dendritic cells (DCreg) in young mice attenuated allogeneic-bone marrow transplant (BMT) induced GVHD, while maintaining the graft versus tumor response ^13-16^. Importantly, our studies also demonstrate amelioration of GVHD in older mice with administration of DCreg following allogeneic BMT [16]. Human DCreg generated from young, healthy donor peripheral monocytes induce regulatory T cells (Treg) and T cell anergy *in* vitro ^17^. These DCreg, however, were generated in fetal bovine serum (FBS)-containing media that potentially exposes the cells to animal immunogens. Additionally, monocytes for DCreg generation must be obtained from HSCT patients prior to transplant, thus requiring cryopreservation to allow for post-transplant treatment with DCreg. To be clinically relevant and for future translational evaluation, the feasibility of generating DCreg in bovine serum-free media from young and older HSCT recipients, as well as the effects of cryopreservation on DCreg generation need to be considered.

## Materials and Methods

### Mononuclear Cell Sources

All samples and informed consent were obtained at the University of Iowa Hospitals and Clinics from “young” (age 30 or below) and “older” (age 50 or above) individuals in accordance with approved IRB-01. A coded identification number was assigned to each sample as obtained. Healthy donor monocytes were purified from de-identified leukoreduction system (LRS) chambers obtained from the DeGowin Blood Center. Peripheral blood (PB) samples were obtained from patients prior to HSCT for varying hematological diseases. These hematological diseases included acute lymphoblastic leukemia (ALL), acute myeloid leukemia (AML), chronic lymphocytic leukemia (CLL), chronic myeloid leukemia (CML), non-Hodgkin lymphoma (NHL), myelodysplastic syndrome (MDS), and myelofibrosis.

### Cell Preparation

Phosphate buffered saline (PBS) was used to flush LRS chambers prior to monocyte enrichment. Prior to transplant, 20-30 mL of whole blood in EDTA was obtained from HSCT recipients and washed with PBS. Following washing, cells from healthy donor and HSCT recipients were labeled with Rosettesep human monocyte enrichment cocktail (Stemcell Technologies, Cambridge, MA) and centrifuged through Histopaque density gradient medium (Sigma-Aldrich, St. Louis, MO). Monocytes were recovered from the interface.

### Cryopreservation

Monocytes obtained from healthy donor LRS cones were cryopreserved at 5 x 10^7^ cells/mL in X-vivo media containing 10% dimethylsulfoxide (DMSO) for 48 hr. in a -80°C freezer followed by transfer into liquid nitrogen until use (3 weeks -12 months).

### DCreg and Conventional Dendritic Cell (cDC) Cultures

Freshly isolated or thawed monocytes were differentiated into DCreg similarly to previously described in serum free media ^17^. Briefly, 1×10^6^ cells/mL were cultured in X-vivo 15 (Lonza, Basel, Switzerland) serum-free media supplemented with human transforming growth factor beta 1 (TGF-β1), granulocyte macrophage-colony stimulating factor (GM-CSF), interleukin 4 (IL-4), and IL-10 (Peprotech, Rocky Hill, NJ; 50 mg/mL each), for 10 days (d.). Human tumor necrosis factor alpha (TNFα; Sigma-Aldrich, St. Louis, MO; 1 µg/mL) was added to the cultures for the last 3 d. of culture. cDC were generated in the presence of only human GM-CSF and IL-4.

### Flow Cytometric Reagents

To prevent non-specific FcγR binding, all cell samples were incubated with anti-CD16/32 (clone 2.4G2) and rat serum. Fluorochrome-conjugated, purified rat immunoglobulins were used as isotype controls for background fluorescence. Cells were stained with the following fluorochrome conjugated monoclonal antibodies (mAbs): CD11c, HLA-DR, CD40, CD80, CD86, PD-L1, ILT-3, CD25, CD103, and Foxp3 (ThermoFisher, Waltham, MA).

### Flow Cytometric Staining and Analysis

Non-adherent DCreg and cDC were recovered from culture and washed in balanced salt solution. Fluorochrome-conjugated or biotinylated mAbs were used to stain cell suspensions. Stained cells were fixed with 0.1% formaldehyde. Flow cytometric data were obtained within 48 hours on a Becton Dickson FACSCanto II or LSR II (San Jose, CA) and analyzed using FlowJo software (FlowJo, LLC, Ashland, OR). Dead cells were excluded via forward/orthogonal light scatter characteristics. Single cells were identified by forward scatter-area versus side scatter-width. Live cells were gated and DC identified as HLA-DR+ CD11c^+^.

### ELISA

At the end of culture, DCreg and cDC culture supernatants were harvested and stored at -80°C until analysis. Supernatants were assayed in triplicate using human/mouse TGFβ1 (Biolegend, San Diego, CA), human IL-6, IL-10, IL-12p70, and TNFα Ready-Set-Go ELISA kits per manufacturers protocols (ThermoFisher, Waltham, MA).

### Treg Induction

As described above, DCreg were generated from donor 1 (Supplementary Fig. 1A). On d. +10 of DCreg culture, CD4+ T cells were purified from donor 2 LRS cone using a Rosettesep human CD4+ enrichment cocktail (Stemcell Technologies, Cambridge, MA; Supplementary Fig. 1B). 5×10^6^ CD4+ T cells from donor 2 were cultured with 5×10^5^ DCreg from donor 1 in X-vivo media for 5 days. Cells were stained with mAbs to CD45RO and Foxp3 to assess the induction of Treg (Supplementary Fig. 1C). Treg were defined as Foxp3+ cells.

**Supplementary Figure 1:**
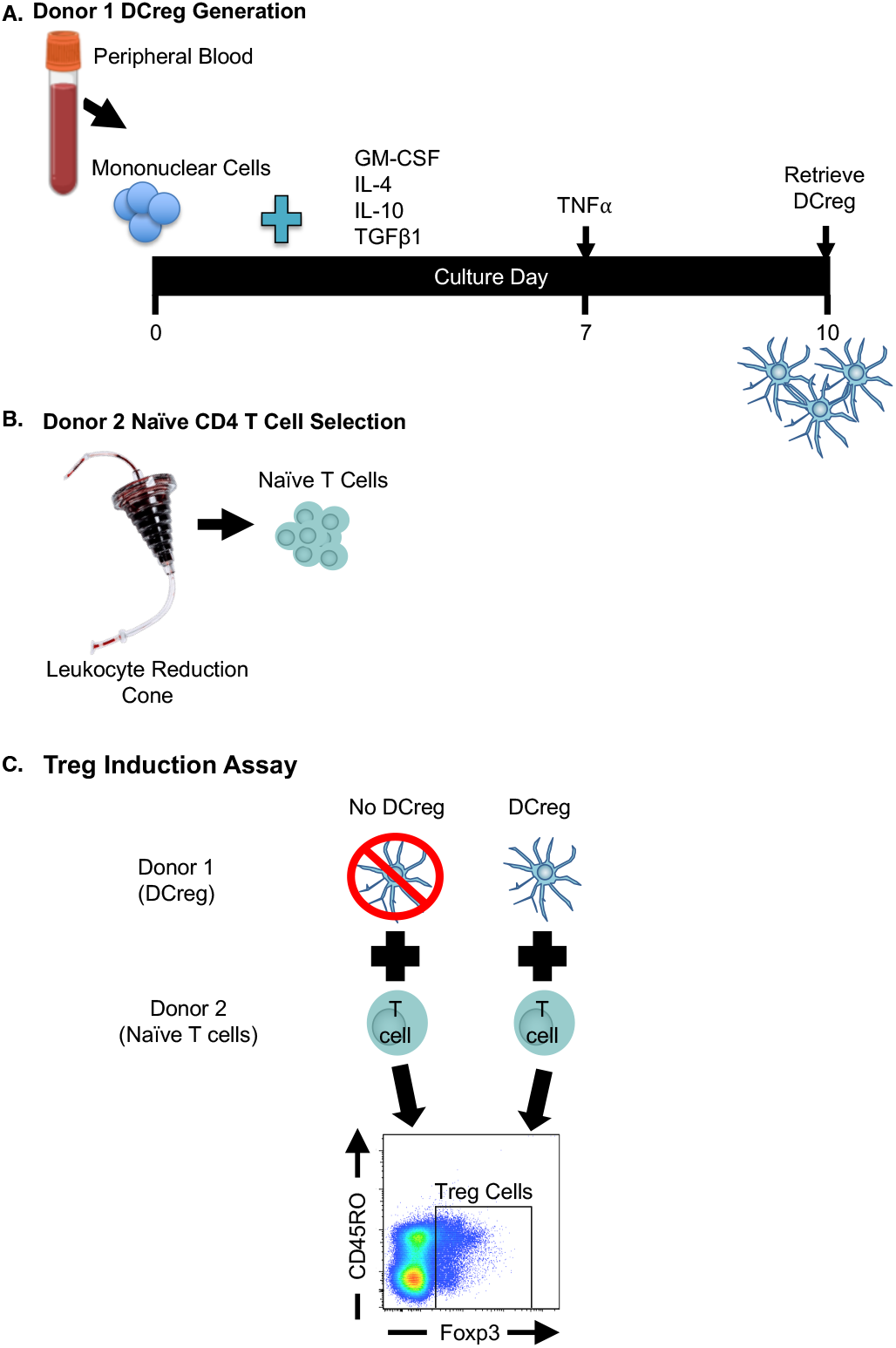
Treg induction Assay. (A) Generation of human DCreg from peripheral blood of donor 1. (B) Naïve donor CD4+ T cells for the Treg induction assay are negatively selected from leukocyte reduction system cone of donor 2. (C) Donor 2 CD4+ T cells are co-cultured without or with Donor 1 DCreg. After 5 days in co-culture, cells are isolated from the culture and stained for CD45RO and intranuclear transcription factor Foxp3. The frequency of Treg is determined via Foxp3+ gating.

### Live cell imaging

Images of patient DCreg were taken at the end of culture with a Nikon TS100 phase contrast inverted microscope equipped with a Retiga 2000R digital camera.

### Statistical Analysis

Statistical significance was determined using a two-tailed Student *t* test or one-way ANOVA and Tukey-Kramer Multiple Comparisons test, where appropriate. The minimal level of confidence deemed statistically significant was *p* value <0.05.

## Results

Because not all HSCT recipients acquire GVHD, the most cost-effective means of treatment would be to generate DCreg on an as needed basis. Thus, monocytes need to be cryopreserved for later Dcreg generation. Unlike murine DCreg ^18^, generation of human DCreg from peripheral monocytes yields a double positive (HLA-DR+ and CD11c+) DC population (Supplementary Fig. 2). This is likely due to a difference in cellular source, as murine DCreg are generated from bone marrow and not peripheral blood. When compared to cDC, young and older DCreg maintained a reduced expression of CD11c and HLA-DR.

**Supplementary Figure 2:**
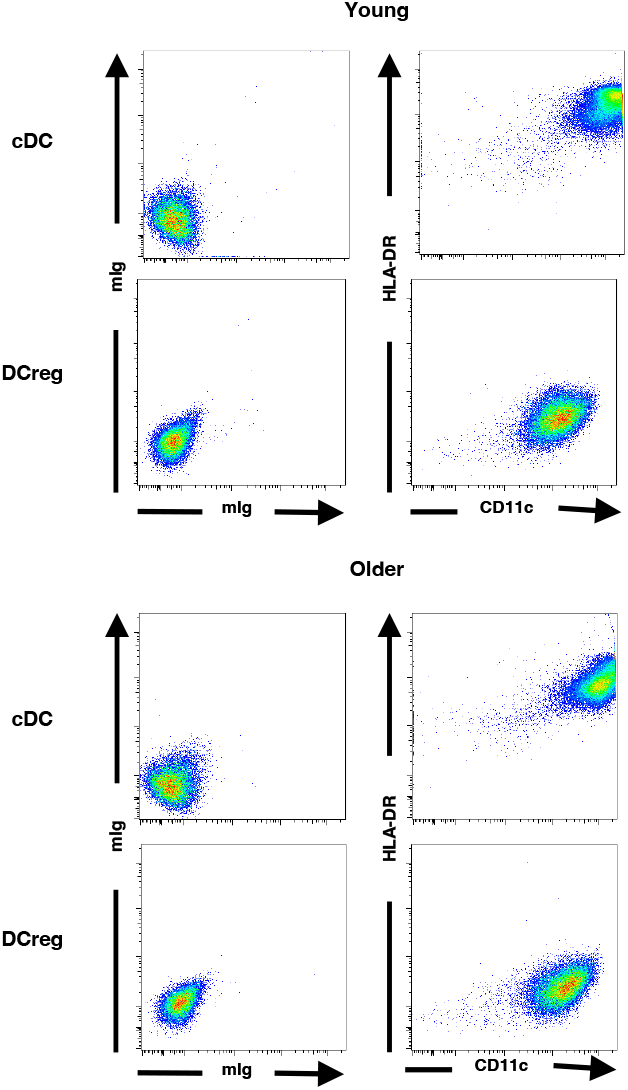
Compared to cDC, young and older human DCreg generated from healthy donors display reduced expression of HLA-DR and CD11c. mIg = isotype control. Representative plots are shown. N ≥ 8 donors per age group.

To assess the effect of cryopreservation, monocytes were allocated into two groups, one for fresh DCreg generation and the other for cryopreservation. For phenotypic analysis and cytokine production evaluation, DCreg were generated from either freshly isolated or cryopreserved monocytes in X-vivo. The number of DCreg generated in culture following cryopreservation was not significantly altered from freshly cultured monocytes (Fig. 1A).

**Figure 1:**
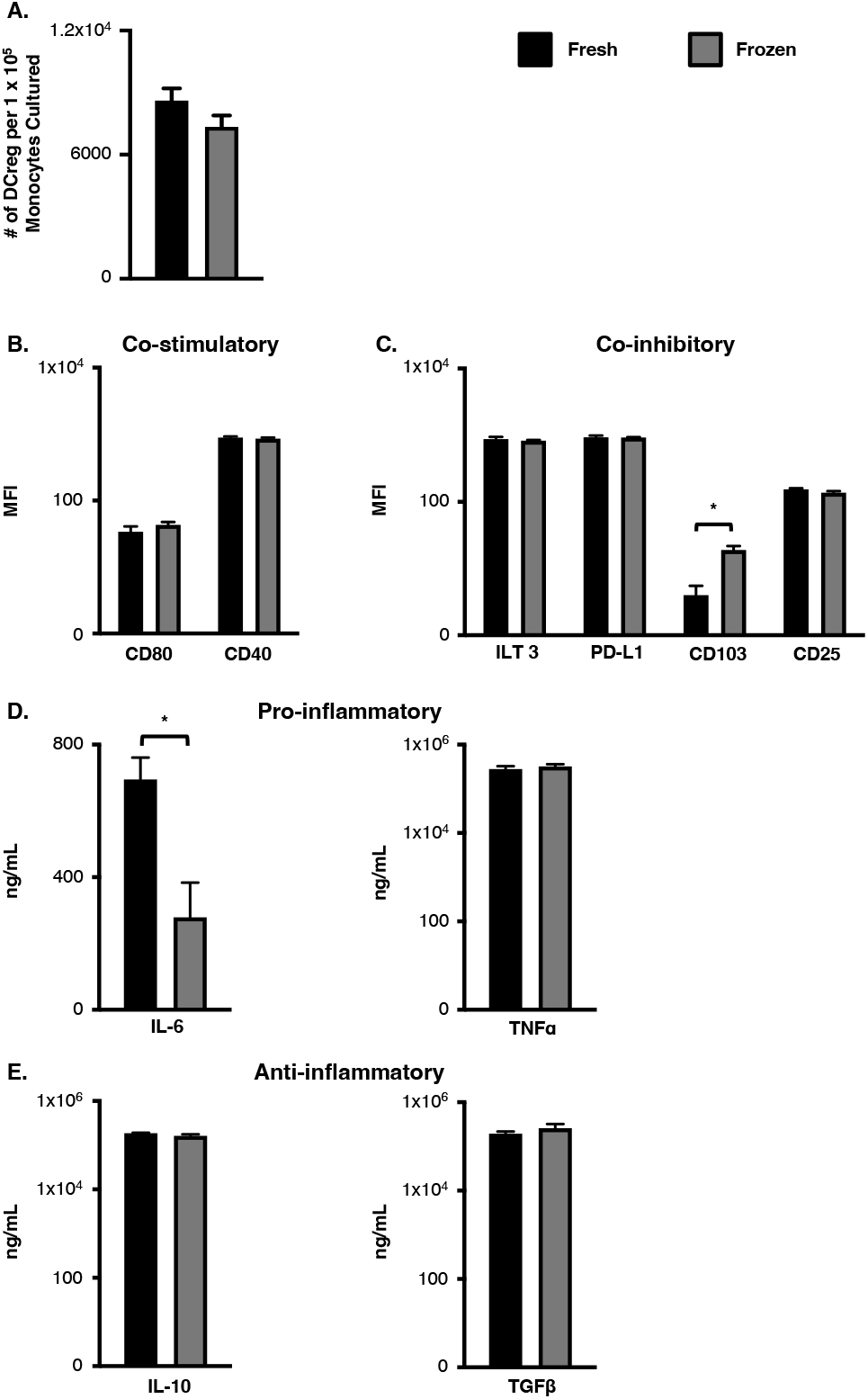
The effects of cryopreservation on DCreg. (A) Freshly isolated and cryopreserved monocytes generated a comparable number of DCreg per 1×10^5^ monocytes cultured. N = 8 per group. Both freshly isolated and cryopreserved monocytes generated DCreg have a similar phenotype that (B) express low levels of co-stimulatory molecules and (C) express high levels of inhibitory molecules. N = 4 per age group. (D) DCreg generated from cryopreserved monocytes secreted less pro-inflammatory IL-6, while (E) production of anti-inflammatory cytokines is maintained. N = 11 per group. Data are mean ± SEM.* = p<0.05.

Furthermore, DCreg generated from cryopreserved monocytes had overall similar levels of co-stimulatory (Fig.1B) and inhibitory molecule (Fig. 1C) expression as DCreg generated from freshly isolated monocytes. The only exception is inhibitory molecule CD103, which was increased on cryopreserved DCreg (Fig. 1C). Secretion of TNFα, IL-10, and TGFβ production from DCreg were comparable between groups (Fig. 1D and E), suggesting DCreg function was not significantly affected by cryopreservation. IL-6 production, however, was significantly reduced following cryopreservation (Fig. 1D). Given overall DCreg surface molecule expression and cytokine production were not altered by cryopreservation, these data support cryopreservation for the generation of DCreg on an as needed basis.

DCreg were generated in X-vivo 15 serum-free media from young and older donor freshly isolated or cryopreserved monocytes and evaluated for phenotype and function. As shown in figure 2A, monocytes from young and older healthy donors generated comparable numbers of DCreg. Phenotypic differences between cDC and DCreg generated from young and older healthy donors was assessed via flow cytometry. As expected, young and older cDC highly expressed the co-stimulatory molecules, CD80 and CD40 (Fig. 2B). Conversely, DCreg had very low expression of CD80 and CD40 compared to cDC, regardless of donor age (Fig. 2B). Additionally, DCreg had significantly higher expression of the inhibitory molecules ILT3, PD-L1, CD103, and CD25 compared to cDC (Fig. 2C). Importantly, older DCreg were phenotypically similar to DCreg generated from young donors (Fig. 2B and C). There was no statistical difference in surface molecule expression between young and older donors on DC populations. The minimal co-stimulatory molecule expression with a high level of inhibitory molecule expression on human DCreg is similar to the phenotype observed in murine DCreg and suggests tolerogenic function.

**Figure 2:**
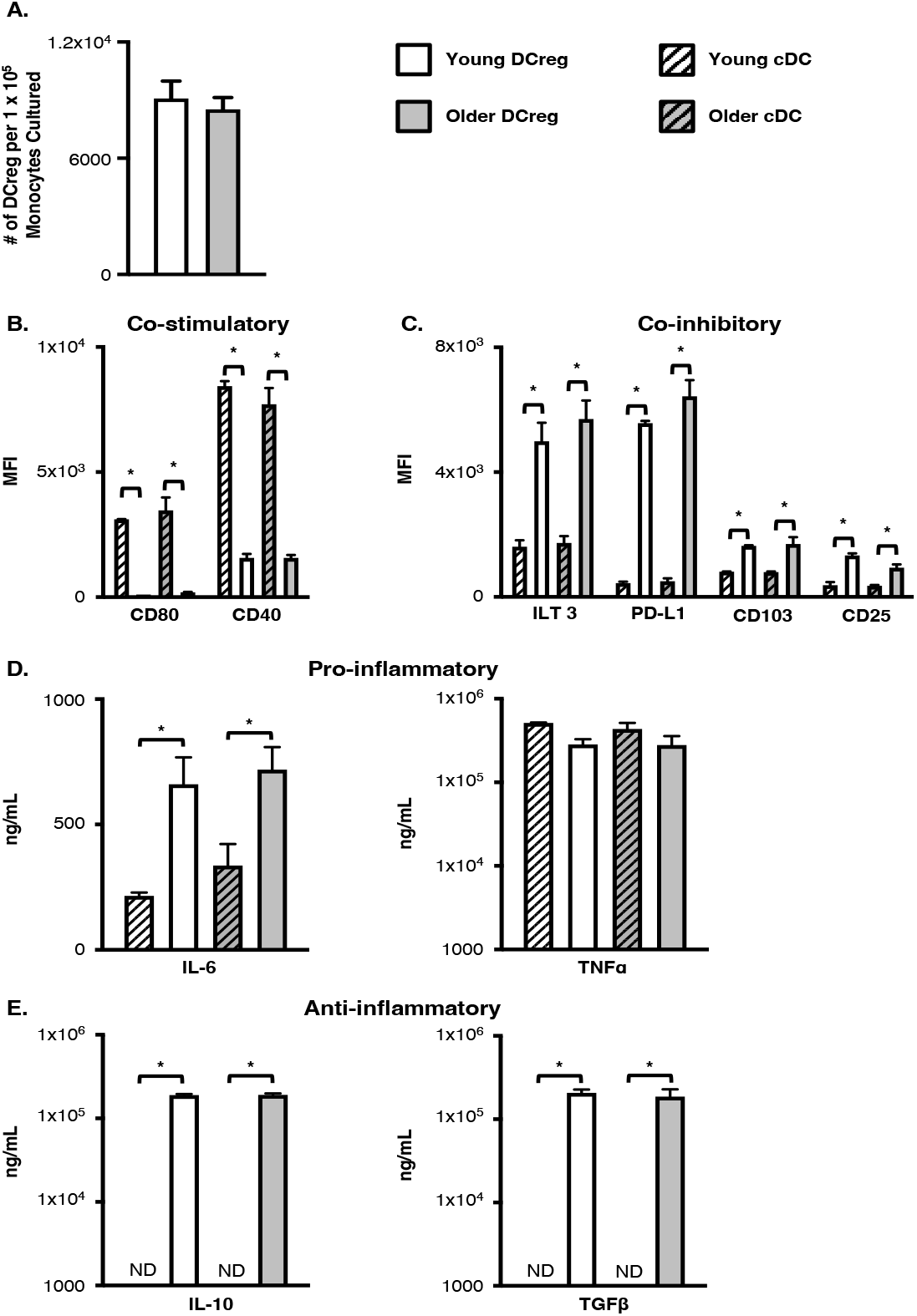
Generation of DCreg in young and older healthy individuals under clinically relevant conditions. (A) DCreg generated from young and older healthy donor monocytes have a similar number of DCreg per 1×10^5^ monocytes cultured. N ≥ 8 donors per age group. Young and older healthy donor monocytes have a similar phenotype that (B) express low levels of co-stimulatory molecules and (C) express high levels of inhibitory molecules. N = 4 per age group. DCreg generated from young and older monocytes secrete similar levels of pro- (D) and anti- (E) inflammatory cytokines. ND, not detected. IL-6, TNFα, and TGFβ ELISAs N = 4 young; N = 4 older. IL-10 production exceeded maximal standard in 1 young and 1 older DCreg culture (exceeded 2 x 10^5^ ng/mL IL-10) and were not included (N = 3 young; N = 3). Data are mean ± SEM. * = p<0.05.

As an indicator of function, cDC and DCreg pro-inflammatory and anti-inflammatory cytokine production was evaluated in culture supernatants at the end of culture. In agreement with previous reports ^19^, cDC, as well as DCreg, generated from monotyes in X-vivo media did not produce IL-12p70 (data not shown). Young and older cDC and DCreg produced comparable amounts of TNFα, while IL-6 production was significantly increased from DCreg (Fig. 2D). Similar to murine DCreg, human DCreg, but not cDC produced significant amounts of IL-10 and TGFβ (Fig. 2E). Production of cytokines was comparable between young and older donors. Interestingly, one young and one older DCreg culture had an IL-10 concentration exceeding 2 x 10^5^ ng/mL and require further dilution to determine the concentration (data not shown).

Hematopoietic disease may negatively impact the generation of DCreg from the monocytes of HSCT patients. In order to address this concern, fifty-six total HSCT patients were enrolled, 12 young and 44 older patients (Supplementary Fig. 3). Acute lymphocytic leukemia (ALL) was the most represented disease in young patients. Acute myeloid leukemia (AML), chronic myeloid leukemia (CML), and non-hodgkin’s lymphoma (NHL) patients were represented in both young and older populations. Although myelodysplastic syndrome (MDS) primarily occurs in older patients, and occurred in 12 of the older patients in our study, MDS was observed in one young patient. Myelofibrosis and chronic lymphocytic leukemia (CLL) patients were only represented in the older population.

**Supplementary Figure 3:**
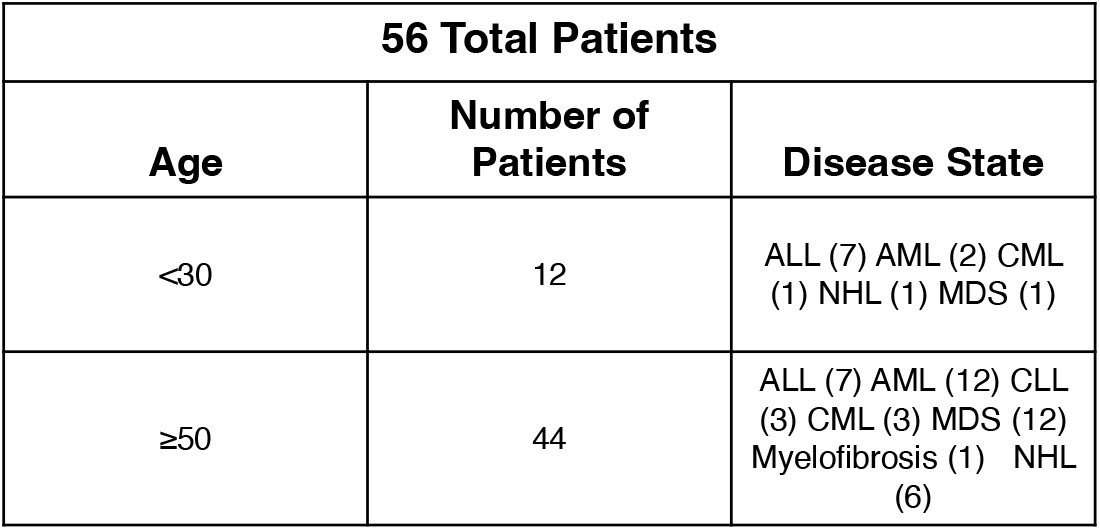
Total number and disease status of young and older patients enrolled in the study.

Similarly to DCreg generated from healthy donors, DCreg generated from young and older HSCT patients expressed HLA-DR and CD11c (Supplementary Fig. 4). Monocytes from young and older HSCT patients generated DCreg, however, older patients’ monocytes produced significantly fewer DCreg (Fig. 3A). Consistent with murine DCreg, young and older patient DCreg expressed similar levels of co-stimulatory (Fig. 3B) and inhibitory molecules (Fig. 3C). Although patient DCreg had low to moderate expression of the co-stimulatory molecules CD80 and CD40 (Fig. 3B), they highly expressed PD-L1 and ILT3 (Fig. 3C). Additionally, patient DCreg expressed CD103 and CD25, although to a lesser extent (Fig. 3C). Production of inflammatory (IL-6 and TNFα) (Fig. 3D) and anti-inflammatory (IL-10 and TGFβ) (Fig. 3E) cytokines were also comparable between young and older patient DCreg. Production of IL-10 was heterogenous between patients. Three of the young and seven of the older patient DCreg had IL-10 concentrations exceeding 2 x 10^5^ ng/mL. As with DCreg generated from healthy donors, patient DCreg did not produce IL-12p70. Monocytes from older MDS patients generated significantly reduced numbers of DCreg compared to non-MDS monocytes (Fig. 3F). Although statistics could not be performed on N=1, monocytes from a young MDS not be performed on N=1, monocytes from a young MDS patient generated fewer DCreg than non-MDS monocytes (Fig. 3F). Interestingly, DCreg cultures from young ALL and AML, as well as older NHL patients, had both uniformly round and elongated spindle shaped cells (Supplementary Fig. 5). Conversely, cultures generated with monocytes from patients with MDS, regardless of treatment regimen, had very few cells and large masses of debris (Supplementary Fig. 5). These findings suggest further investigation into better methods for generating DCreg from MDS patients is required. It is possible MDS patients are not good candidates for DCreg treatment due to the inability to efficiently generate DCreg from these individuals.

**Supplementary Figure 4:**
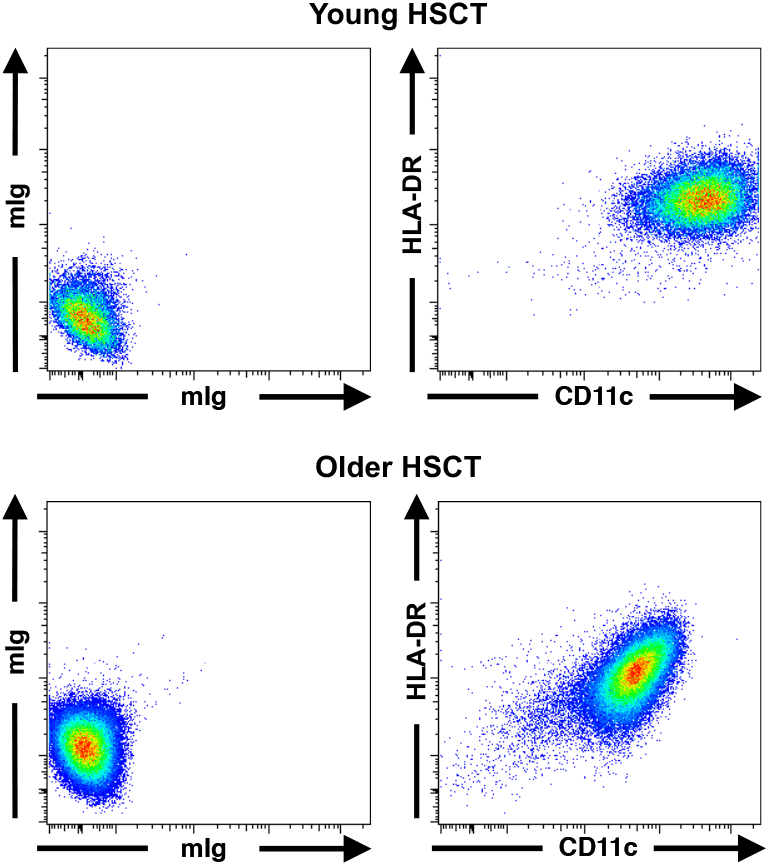
Young and older HSCT patient generated DCreg express low levels of HLA-DR and CD11c. DCreg were generated from young and older monocytes isolated from the peripheral blood of patients prior to HSCT . mIg = isotype control. Representative plots are shown. Young N = 10; Older N = 23.

**Figure 3:**
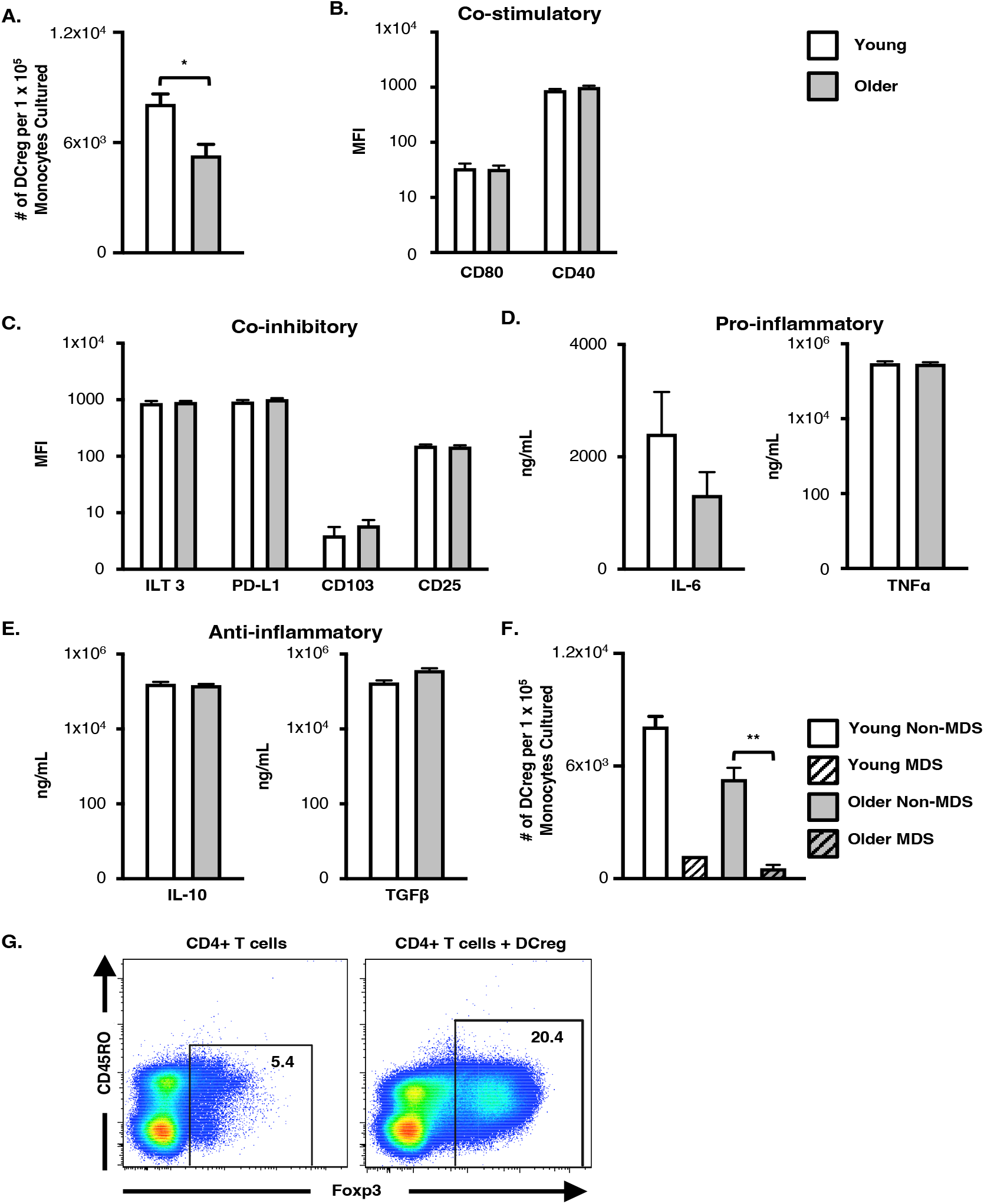
Generation of DCreg from young and older patients just prior to HSCT. (A) Number of DCreg generated per 1 x 10^5^ monocytes cultured in yound and older HSCT patients. Samples that did not grow were excluded. Young N = 11; Older N = 26. HSCT DCreg expression of (B) -stimulatory molecules and (C) co-inhibitory molecules. Young N = 10; Older N = 23. (D) Pro- and anti- (E) inflammatory cytokine production by DCreg generated from young and older HSCT patients. IL-6, TNFα, and TGFβ ELISAs N = 7 young; N = 16 older. IL-10 production exceeded maximal standard in 3 young and 7 older DCreg culture (exceeded 2 x 10^5^ ng/mL IL-10) and were not included (N = 4 young; N = 9). (F) Young and older MDS patient DCreg numbers. Young non-MDS N = 11; Young MDS N = 1; Older non-MDS N = 24; Older MDS N = 7. Statistical comparison between young and older MDS DCreg numbers was not performed due to low sample size. ** = p<0.01. (G) DCreg generated from healthy monocytes induce Foxp3+ Treg. Representative plots are shown. N = 3. Data in A-F are mean ± SEM. * = p<0.05.

**Supplementary Figure 5:**
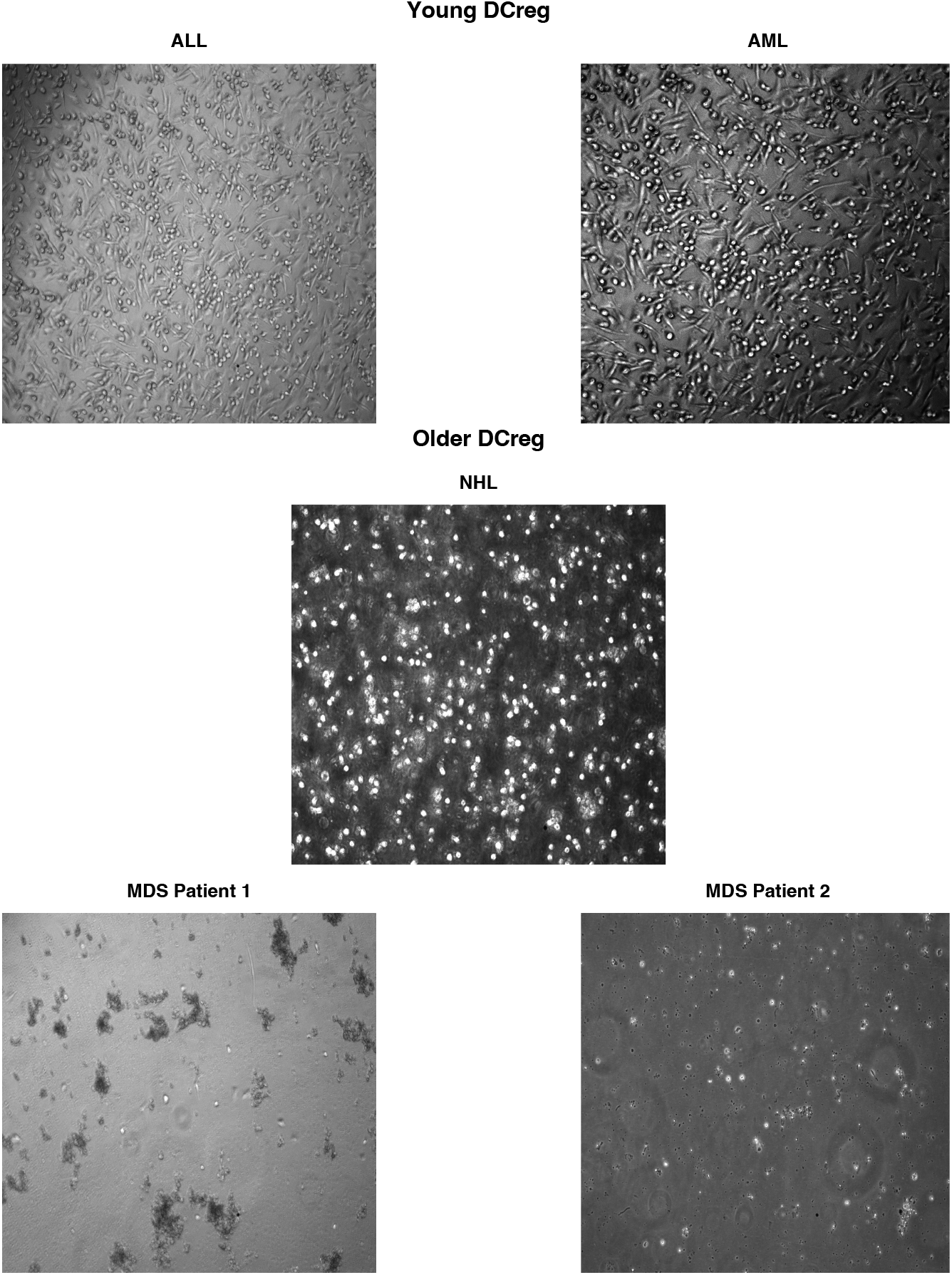
Size and shape of DCreg generated from HSCT patients. DCreg generated from monocytes purified from ALL, AML, and NHL patients showed round and elongated spindle shaped cells while DCreg generated from MDS patients were irregular in shape with large masses of debri. Live cell images (40X) using a phase contrast inverted microscope were taken at the end of the culture period. MDS images are representative of MDS patients. The ALL, AML, and NHL images are representative of the ALL, AML, CML, and NHL (non-MDS) patients in the study. Young non-MDS N = 11. Young MDS N = 1. Older non-MDS N = 24. Older MDS N = 7.

DCreg function was further confirmed through the ability of DCreg to induce Treg. On d. +10 of DCreg culture (donor 1), naïve CD4+ T cells were purified using a CD4+ Rosettesep enrichment cocktail from a second LRS chamber (donor 2) (Supplementary Fig. 1A & 1B). CD4+ T cells were either placed in culture alone or put into culture with DCreg at a 10:1 ratio for 5 d., and then stained for Foxp3 via intranuclear staining. Approximately 5% of the CD4+ T cells were Foxp3+ at baseline, as indicated in the T cell only cultures (Fig. 3G). Confirming DCreg function in cells generated in serum-free media, a higher frequency of Treg cells were present in cultures containing DCreg compared to T cell only cultures (Fig. 3G). Additionally, this equated to a numerical difference of approximately 5 x 10^4^ Treg per 1 ×10^6^ T cells cultured, compared to 2 x 10^4^ Treg per 1 x 10^6^ T cells in the T cell only cultures. Taken together, these data indicate DCreg generated from young and older healthy donors, in serum-free media, are phenotypically and functionally tolerogenic.

## Discussion

DCreg generated from young, syngeneic BM has been demonstrated to alleviate GVHD-induced mortality in young mice; importantly, the GVT response remained intact following DCreg treatment ^13-16^. We previously demonstrated that DCreg generated from older mice reduce GVHD-associated morbidity and mortality in older BMT recipient mice ^18^. Recent work in rats has shown that DCreg alleviate autoimmune disease via formation of microchimerism in treated rats ^20^. These data provide support for the potential of DCreg therapy in the long-term amelioration of human GVHD. Previous studies have demonstrated DC with low co-stimulatory molecule expression and the ability to induce Treg and T cell anergy *in vitro* can be generated from young, healthy donor monocytes in FBS-containing media ^17^. Further, treatment with anti-inflammatory agents or immunosuppressive drugs has been shown to result in DCreg differentiation ^21^. The use of animal serum is not compliant with good manufacturing practices (GMPs) and, to our knowledge DCreg generation from older individuals or from HSCT patients have not been addressed. Also, to be clinically relevant and economically feasible, monocytes would need to be cryopreserved for generation of DCreg on a case-by-case basis.

Cryopreserved monocytes from young and older healthy donors generated similar numbers of DCreg as freshly isolated monocytes. DCreg generated from cryopreserved monocytes maintained high expression of inhibitory molecules and anti-inflammatory cytokine production. There were no age-dependent variations between cryopreserved and freshly isolated monocytes. These data suggest DCreg generated from cryopreserved monocytes are phenotypically and functionally similar to those generated from freshly isolated monoctes. Further confirmation of function may be evaluated via induction T cell anergy in future studies.

This study demonstrates DCreg can be generated in serum free media from young and older healthy donor and HSCT patient monocytes. Although DCreg generation from young and older healthy monocytes generated similar numbers of DCreg, older patient monocytes generated fewer DCreg per monocyte cultured compared to young patient monocytes.

In agreement with DCreg generated in serum-containing media ^17^, DCreg generated in X-vivo serum-free media from healthy donors and HSCT patients maintained low co-stimulatory molecule expression. CD25, CD103, PD-L1, and ILT3 have been reported on various DC populations and are thought to play a role in regulating immune activation and tolerance ^20,22-33^. Human DC that inhibited T cell function were shown to co-express CD25 and IDO ^32^. Indeed, all human DCreg generated in this study expressed CD25. As murine DCreg also produced IDO and were shown to be necessary for maximal protection from GVHD, it would be of interest to evaluate human DCreg IDO expression. We have previously demonstrated PD-L1 and PIR B expression on DCreg is required for protection from GVHD in BMT mice ^18^. Human healthy donor and patient DCreg generated in X-vivo media also highly expressed PD-L1 and ILT3.

IL-10 and TGFβ are anti-inflammatory cytokines involved in Treg function; TGFβ has also been shown to be critical in the development and maintenance of Treg ^34,35^. In accordance with previous reports describing DC with regulatory functions ^28,36-38^ and our murine DCreg ^18^, DCreg generated *in* vitro from healthy and HSCT patient monocytes produced substantial amounts of IL-10 and TGFβ. Confirming human DCreg function and in accordance with previous reports describing DCreg that induce Treg and T cell anergy ^17,28,36,38-42^, human DCreg generated in X-vivo induced Treg. Induction of T cell anergy by human DCreg was not evaluated in the current study. Although the requirement for specific inhibitory molecules and/or anti-inflammatory cytokines was outside the scope of the current study, the phenotypic and functional data suggest human DCreg and murine DCreg may function through similar mechanisms.

For the first time, these data demonstrate DCreg generated under clinically relevant conditions (i.e. in serum-free media and from cryopreserved monocytes) have a similar phenotype and function as DCreg generated from freshly isolated monocytes. Additionally, DCreg can be generated from older healthy donors with similar numbers, phenotype, and function to young donors. Perhaps most importantly, DCreg can be reliably generated from young and older HSCT patients with a wide variety of hematological diseases. These DCreg express comparable inhibitory surface molecules and produce anti-inflammatory cytokines as DCreg generated from healthy donor monocytes. Many of the DCreg cultures from MDS patients (5/12 older patients and 1/1 young patient) had considerable debris and very low cell recovery, preventing phenotypic analysis and cellular yield inclusion. These findings suggest further investigation into alternative methods for generating DCreg from MDS patients is required. It is possible MDS patients are not good candidates for DCreg treatment due to the inherent defects in monocytes resulting in an inability to efficiently generate DCreg from these individuals.

## Conclussion

Our study demonstrates DCreg can be generated under clinically relevant conditions from both young and older individuals. Further, to increase personalization and feasibility, DCreg can be generated following cryopreservation of monocytes on an as needed basis. Lasty, DCreg generated from the peripheral blood of HSCT patients prior to transplant are phenotypically and functionally active. Collectively, our study provides critical data to support the use of *in vitro* generated DCreg as a therapy to reduce GVHD-associated morbidity and morbidity in HSCT recipients.

## Acknowledgements

The authors wish to thank the Carver College of Medicine Flow Cytometry Core for their assistance in collecting flow data, Karen Parrott, RN for identifying and consenting patients, Doug Scroggins for experimental assistance, Alexis Potratz for formatting assistance, and Jackie Bickenbach, PhD for assistance with live cell imaging.

